# Host lipid alterations after *Macrophomina phaseolina* infection contribute to charcoal rot disease susceptibility in grain sorghum: Evidence from transcriptomic and lipidomic data

**DOI:** 10.1101/854067

**Authors:** Ananda Y. Bandara, Dilooshi K. Weerasooriya, Sanzhen Liu, Christopher R. Little

**Author notes:** Department of Plant Pathology and Environmental Microbiology, 211 Buckhout Lab, Pennsylvania State University, University Park, PA 16802. Corresponding author: Christopher R. Little.

## Abstract

- Lipids are involved in central metabolic processes and confer basic configuration to cellular and subcellular membranes. Lipids also play a role in determining the outcome of plant-pathogen interactions. Lipid based links that delineate either host resistance or susceptibility against necrotrophic microorganisms are poorly investigated and described. *Macrophomina phaseolina* (MP) is an important necrotrophic fungus which causes diseases in over 500 plant species including charcoal rot in sorghum.
- We used RNA sequencing and automated direct infusion electrospray ionization-triple quadrupole mass spectrometry (ESI-MS/MS) to quantitatively profile the transcriptomes and lipidomes of a known charcoal rot resistant (SC599) and susceptible (Tx7000) sorghum genotype in response to MP inoculation.
- We found that MP is capable of significantly decreasing the phosphatidylserine, phytosterol, and ox-lipid contents in the susceptible genotype while significantly increasing its stigmasterol:sitosterol and monogalactosyldiacylglycerol: digalactosyldiacylglycerol ratios. None of the above was significantly affected in the resistant genotype, except for the significantly increased ox-lipid content.
- Our transcriptome and functional lipidome findings suggested the lethal impacts of MP inoculation on plastid- and cell- membrane integrity and the lipid based signaling capacity of the charcoal rot susceptible sorghum genotype, Tx7000. Findings also suggested the strong oxidative stress experienced by Tx7000 under MP inoculation and shed light on the potential lipid classes involved in induced charcoal rot disease susceptibility.

## Introduction

Lipids play important and indispensable roles in many physiological processes in living organisms. They are involved in central metabolism and confer basic configuration to cell and organelle membranes. Membranes are fundamental to cell structure and function. Therefore, maintenance of membrane integrity and fluidity is required for plants to survive under environmental changes (Wallis & Browse, 2002; Welti *et al*., 2007). The membrane trafficking, exo- and endocytosis, cytoskeletal rearrangements, photosynthesis, and signal transduction are some of the key functions played by lipids in eukaryotes (Wang, 2004; van Leeuwen *et al*., 2004; Funk, 2001; Shea & Del Poeta, 2006).

Lipids are significant determinants of plant-pathogen interactions. For instance, preformed structural barriers such as the cuticle contribute to first line defense against plant pathogens (Reina-Pinto & Yephremov, 2009). Cutin, a polyester of hydroxy and epoxy-hydroxy C16 and C18 fatty acids, is the major constituent of the cuticle (Kolattukudy, 2001), which provides a physical barrier between the pathogen and host cell (Jenks *et al*., 1994). The cell membrane is also a structural barrier for pathogen entrance. Phospholipids such as phosphatidylcholine (PC), phosphatidylethanolamine (PE), phosphatidylinositol (PI), phosphatidylglycerol (PG), phosphatidylserine (PS) are the important lipid constituents that make cell and mitochondria membranes (Horvath & Daum, 2013) while the galactolipids, monogalactosyldiacylglycerol (MGDG) and digalactosyldiacylglycerol (DGDG), are the major lipid constituents of chloroplast membranes (Joyard *et al*., 2010; Boudière *et al*., 2014; Fujii *et al*., 2014). Phytosterols are also constituents of plant cell membranes. Other than their structural contribution, phytosterols have also been shown to play a crucial role in plant innate immunity against phytopathogens (Wang *et al*., 2012).

Lipids and fatty acids are also important as signaling molecules in plant defense against phytopathogens (Walley *et al*., 2013; Kachroo & Kachroo, 2009; Shah, 2005; Laxalt & Munnik, 2002). Polyunsaturated fatty acids such as linoleic acid can enzymatically or non-enzymatically be oxygenated to produce oxylipins, which have diverse signaling properties in mammals, microbes, and plants (Walley *et al*., 2013; Kachroo & Kachroo, 2009; Shea & Del Poeta, 2006; Howe, 2007). Jasmonic acid is an extensively studied oxylipin in plants. Moreover, mechanical, biotic, and low-temperature stresses increase many membrane lipids with oxidized acyl chains (i.e. ox-lipids) in *Arabidopsis thaliana* (Vu *et al*., 2012, 2014). Ox-lipids may be produced enzymatically through the action of lipoxygenase or non-enzymatically through the action of reactive oxygen species (ROS) (Zoeller *et al*., 2012). Like oxylipins, ox-lipids also function as signaling molecules that initiate stress responses in plants (Andersson *et al*., 2006). Phosphatidic acid (PA) (a phospholipid) is a potent signaling molecule in plants and animals that modulates the activities of kinases, phosphatases, phospholipases, and proteins involved in membrane trafficking, Ca2+ signaling, and the oxidative burst (Munnik, 2001; Wang, 2004). The role of PA in plant defense against pathogens is well documented (Laxalt & Munnik, 2002; Munnik, 2001; Wang, 2004). Lipids play diverse and pivotal roles in determining the outcome of plant-pathogen interactions.

*Macrophomina phaseolina* is a soil-borne necrotrophic fungal pathogen and is reported to causes diseases in over 500 plant species (Islam *et al*., 2012). Despite its broad host range, *Macrophomina* is a monotypic genus and contains only one species: *M. phaseolina* (Sutton, 1980). It can remain viable in soil and crop residue for more than four years (Short *et al*., 1980). Higher temperatures (30-35°C) and low soil moisture are conducive for the diseases caused by *M*. *phaseolina* including seedling blight, charcoal rot, stem rot, and root rot (Sandhu *et al*., 1999). Therefore, drought-prone regions are highly vulnerable to *M*. *phaseolina-*associated crop losses. Increased occurrence of the pathogen on various crop species has been recently reported worldwide (Khangura & Aberra, 2009; Mahmoud & Budak, 2011).

*M*. *phaseolina* causes charcoal rot disease in many economically important crops such as sorghum, soybean, maize, alfalfa and jute (Islam *et al*., 2012). Charcoal rot is a high priority fungal disease in sorghum [*Sorghum bicolor* (L.) Moench], causing tremendous crop losses wherever sorghum is grown (Tarr, 1962, Tesso *et al*., 2012). Recent studies have shown that charcoal rot can negatively affect the grain sorghum physical and chemical properties (Bandara et al., 2017a), yield parameters (Bandara et al., 2017b), and leaf greenness (Bandara et al., 2016), as well as the key biofuel traits of sweet sorghum (Bandara et al., 2017c). There are limited options available to control charcoal rot disease. Moreover, the underlying molecular mechanism/s of charcoal rot resistance is/are currently unknown. Therefore, there is a pressing need to understand the molecular basis of charcoal rot resistance in sorghum before resistance is deployed as an effective control strategy. The genetic control of resistance to necrotrophic pathogens in general and *M*. *phaseolina*, in particular, is poorly understood and large scale gene expression studies and complimentary functional studies such as lipidomics assays can provide a broader view and better understanding on the disease resistance mechanisms. Here we investigate the stalk tissue transcriptomes of known charcoal rot resistant and susceptible sorghum genotypes after challenged with *M. phaseolina* using Illumina sequencing technology. Furthermore, we take advantage of automated direct infusion electrospray ionization-triple quadrupole mass spectrometry (ESI-MS/MS) to quantitatively profile the lipidome from sorghum stalk tissues after *M. phaseolina* infection. Plant lipidomics based on ESI-MS/MS is a useful method to study the responses of hundreds of lipid molecular species to various environmental stresses (Zheng *et al*., 2011; Welti *et al*., 2002). Therefore, the objectives of the current study were to (i) identify differentially expressed lipid metabolism related genes between charcoal rot resistant and susceptible sorghum genotypes in response to *M. phaseolina* inoculation using RNA-Seq, (ii) identify the differentially expressed lipid classes and species between charcoal rot resistant and susceptible sorghum genotypes in response to *M. phaseolina* inoculation and (iii) uncover the potential links between lipids and charcoal rot resistance or susceptibility at the transcriptional and functional lipidomics levels.

## Materials and Methods

### Plant materials, establishment and maintenance, inoculum preparation and inoculation

Two experiments (each arranged in completely randomized design) were conducted in greenhouse facilities at the Kansas State University, USA in 2013 and 2016 to obtain stalk tissues for RNA sequencing and lipodomics investigations, respectively. For both experiments, commonly used charcoal rot resistant (SC599R) and susceptible (Tx7000) sorghum genotypes were used. In both experiments, the fungicide, Captan (N-trychloromethyl thio-4-cyclohexane-1,2 dicarboxamide) treated seeds were planted (3 seeds per pot) in 19 L Poly Tainer pots filled with Metro-Mix 360 growing medium (Sun Gro Bellevue, WA, U.S.A). At three weeks after emergence, each pot was thinned to a single vigorous seedling. Seedlings/plants husbandry was carried out according to the procedures described by Bandara *et al*., (2015). For both experiments, plants were maintained under a 16-h light/8-h dark photoperiod at 25-32°C. The *M. phaseolina* isolate used for this study was obtained from the row crops pathology lab at the Department of Plant Pathology, Kansas State University and was previously characterized to be a highly virulent isolate. Inoculum was prepared following the protocol described by Bandara *et al*., (2015). Briefly, *M. phaseolina* was grown on potato dextrose broth to obtain *M. phaseolina* mycelia. The broth containing mycelial mass was blended and filtered to obtain small hyphal fragments. Filtrates containing hyphal fragments were centrifuged and the resulting pellets were resuspended in sterile phosphate-buffered saline (PBS) and concentration was adjusted to 2 × 10^6^ hyphal fragments mL^-1^ by adding appropriate amount of PBS. All inoculum preparation steps were performed under aseptic conditions. Plants were inoculated through injection method at 14 days after anthesis by injecting (1 mL sterile surgical syringe, 1.5 inch/26 gauge needle) 0.1 mL of inoculum into the basal internode of the stalk. PBS (pH 7.2) was used as the mock-inoculated control treatment.

## RNA sequencing

### Tissue collection, RNA extraction and quality check, cDNA library preparation and Illumina sequencing

Stalk samples were collected at 2, 7, and 30 days post inoculation (DPI) from three biological replicates after receiving the *M. phaseolina* and mock-inoculation treatments (3 biological replicate per DPI per treatment per sorghum line = 36 plants altogether). During sample collection, approximately 8-10 cm long stalk piece encompassing the inoculation point was cut off from each plant and immediately dipped in liquid nitrogen to avoid mRNA deterioration. Stalk pieces were then stored at −80°C until RNA extraction. For RNA extraction, approximately 1 g of stalk tissues 1 cm above the symptomatic region was chopped in to liquid nitrogen and ground in to a fine powder using mortar and pestle. Following the manufacturer’s instructions, total RNA was extracted using Triazole reagent (Thermo Scientific, USA). RNA was subsequently treated with Amplification Grade DNAse I (Invitrogen Corporation, USA). Using Nanodrop 2000 instrument (Thermo Scientific, USA), RNA quality and quantity was determined. Samples were diluted with appropriate volume of RNase free water to adjust the RNA concentration to recommended range (100-200ng/ul). Lastly, the RNA integrity and quantity of diluted samples were checked using Agilent 2100 Bioanalyzer (Agilent Technologies Genomics, USA) before samples were used for cDNA library preparation. Following the manufacturer’s protocol (Illumina Inc., USA), 36 cDNA libraries were constructed using Illumina TruSeq^TM^ RNA sample preparation kit. Using “oligodT” attached magnetic beads, each RNA sample was enriched (twice) for poly-A mRNAs. Following the manufacturer’s protocol, (Illumina Inc., USA), purified mRNA was chemically fragmented and subsequently converted to single-stranded cDNA. Each cDNA library was distinctly barcoded with unique adapter indexes. Libraries were sequenced using HiSeq 2000 platform (Illumina Inc., USA) with 100bp single-end sequencing runs at the Kansas University Medical Center Genome Sequencing Facility.

### Sequence reads processing, mapping to sorghum reference genome, analysis for differentially expressed genes and assigning gene functions

Sequencing reads were subjected to adapter trimming and quality filtering using an adapter trimmer “Cutadapt” (Martin, 2011). Reads were then aligned to the *sorghum bicolor* reference genome (Sbicolor_v1.4) (Paterson *et al*., 2009) using Genomic Short-read Nucleotide Alignment Program (GSNAP) (Wu & Watanabe, 2005). Differential gene expression analysis was performed using ‘DESeq2’ package. Analysis was conducted for each informative gene within a given post inoculation stage (2, 7, and 30 DPI). The genes with significant genotype × inoculation treatment interaction (genotype is with 2 levels: resistant and susceptible sorghum lines; inoculation treatment is with 2 levels: infected with *M. phaseolina* and mock-inoculated control) were considered as significantly differentially expressed. In order to control the inflation of type I error due to simultaneous inferences (multiple comparisons), 5% false discovery rate (FDR) was deployed. For this, a q-value (Benjamini and Hochberg, 1995) was computed for each gene and only the genes with q-values < 0.05 were considered as significantly differentially expressed (i.e. significant two way interaction). The differentially expressed genes were functionally annotated using the “Phytozome” data base (Goodstein et al., 2012). The associated metabolic pathways of differentially expressed genes were determined using SorghumCyc data base (http://pathway.gramene.org/gramene/sorghumcyc.shtml). Finally, the metabolic pathway enrichment analysis was conducted according to the procedure described by Dugas et al. (2011) to identify the significantly enriched metabolic pathways.

### Lipidome analysis

#### Lipid extraction

Stalk tissues were collected from *M. phaseolina* inoculated and mock-inoculated control plants from resistant and susceptible sorghum genotypes at 4, 7, and 10 days post-inoculation (DPI) and used for lipid extraction (five biological replicates per DPI per treatment per sorghum line = 60 plants altogether). At sampling, approximately 1 g of stalk harvested 1 cm away from the symptomatic area was chopped in to 6 mL of isopropanol with 0.01% butylated hydroxytoluene (BHT) [preheated to 75°C] in a 50 mL glass tube with a Teflon lined screw-cap (Thermo Fisher Scientific, Inc., Waltham, MA, USA). Tubes were incubated in a waterbath at 75°C for 15 min to inactivate lipid-hydrolyzing enzymes. After reaching room temperature, 3 mL of chloroform and 1.2 mL of water were added to each tube and stored at −80°C until further processing. The lipid extraction was performed following the protocol described by Vu *et al*., (2012). Briefly, the lipid extract in isopropanol, BHT, chloroform and water was shaken on an orbital shaker at room temperature for 1 h and transferred to a new glass tube using a Pasteur pipette, leaving the stalk pieces in the original tube. Subsequently, 8 mL of chloroform: methanol (2:1) mixture was added to the stalk pieces and shaken on an orbital shaker (140 rpm) at room temperature for 1 h. The resulting solvent was transferred to the first extract. The addition, shaking and transfer steps were carried out four times including one overnight shake until the stalk pieces of every sample became white. Then the solvent was evaporated in an N-EVAP 112 nitrogen evaporator (Organomation Associates, Inc., Berlin, MA, USA), leaving the lipid extract. Lastly, the lipid extract was dissolved in 1 mL of chloroform and stored at −80 °C. The remaining stalk pieces of each sample were dried overnight in an oven at 105°C, cooled and weighed to express the lipid content on a dry weight basis. Dry weights were measured using a balance (Mettler Toledo AX, Mettler Toledo International, Inc., Columbus, OH, USA) with a detection limit of 2 μg.

### Lipid profiling with electrospray ionization-triple quadrupole mass spectrometer

Automated electrospray ionization-tandem mass spectrometry approach was used for lipid profiling. Data acquisition, analysis, and acyl group identification were performed following the methods described by Xiao *et al*. (2010) with modifications and an added quality-control approach. From the lipid extracts that were dissolved in 1 mL of chloroform, an aliquot of 15 to 70 μL (corresponding to approximately 0.2 mg dry weight) was added to each of two vials (vial 1 and vial 2). Following the methods described by Welti *et al*., (2002), accurate amounts of internal standards were measured and added to vial 1 in the following quantities: 0.6 nmol phosphatidylcholine (PC) (di12:0), 0.6 nmol PC (di24:1), 0.6 nmol lysophosphatidylcholine (LPC) (13:0), 0.6 nmol LPC (19:0), 0.3 nmol phosphatidylethanolamine (PE) (di12:0), 0.3 nmol PE (di23:0), 0.3 nmol lysophosphatidylethanolamine (LPE) (14:0), 0.3 nmol LPE (18:0), 0.3 nmol phosphatidylglycerol (PG) (di14:0), 0.3 nmol PG (di20:0(phytanoyl)), 0.3 nmol lysophosphatidylglycerol (LPG) (14:0), 0.3 nmol LPG (18:0), 0.23 nmol phosphatidylinositol (PI) (16:0–18:0), 0.16 nmol PI (di18:0), 0.2 nmol phosphatidylserine (PS) (di14:0), 0.2 nmol PS (di20:0(phytanoyl)), 0.3 nmol phosphatidic acid (PA) (di14:0), 0.3 nmol PA (di20:0(phytanoyl)), 0.31 nmol TAG (tri17:1), 0.36 nmol digalactosyldiacylglycerol (DGDG) (16:0–18:0), 0.95 nmol DGDG (di18:0), 1.51 nmol monogalactosyldiacylglycerol (MGDG) (16:0–18:0) and 1.3 nmol MGDG (di18:0). Only the last four internal standards were added to vial 2, in half the amount as vial 1. The solvents [chloroform: methanol: 300 mM ammonium acetate in water, 300:665:35 (v/v/v)] were added to the lipid extract and internal standard mixture in each vial. The final volume was 1.4 mL. Unfractionated lipid extracts were introduced by continuous infusion into the electrospray ionization (ESI) source on a triple quadrupole MS/MS (API4000, ABSciex, Framingham, MA, USA) using an autosampler (LCMini PAL, CTC Analytics AG, Zwingen, Switzerland) at 30 μL min^-1^. Data and spectra acquisition, resolution adjustment of mass analyzers, background subtraction from each spectrum, data smoothening and peak area integration, data processing, and calculation of normalized lipid intensities were performed according to the procedures described by Vu *et al*. (2014). The lipid values are reported as normalized intensity (%) per mg stalk dry weight, where a value of one is the intensity of 1 nmol of internal standard.

### Statistical analysis of lipid data

Lipid data were analyzed for variance (ANOVA) using the PROC GLIMMIX procedure of SAS software version 9.2 (SAS Institute, 2008). Analyses were conducted at lipid class level (DGDG, MGDG, SQDG, PG, phosphatidylcholine, PE, PI, PS, PA, lysoPC, lysoPE, sterol glucosides, acyl(18:2) sterol glucosides, acyl(16:0) sterol glucosides, NL297(18:2) containing TAG, NL295(18:3) containing DAG/TAG, NL273(16:0) containing DAG/TAG, HexCer, prec291(18:3-2O) or 18:4-O, prec293(18:2-2O) or 18:3-O, and ratios of MGDG:DGDG, PE:PC, galactolipids [DGDG, MGDG, SQDG]/phospholipids [PG + PC + PE + PI + PS + PA]) and individual lipid species levels. Although 227 different lipid species were detected, based on the limit of detection (>0.002 nmol) and coefficient of variation (< 0.3) criteria for pooled samples, only 132 were qualified for the final ANOVA analysis (see supplementary table 1). The restricted maximum likelihood (REML) method was used to estimate variance components. Genotype (SC599, Tx7000), inoculation treatment (*M. phaseolina*, control) and time point (4, 7, and 10 DPI) were considered fixed factors. Model assumptions were tested using studentized residual plots (for identical and independent distribution of residuals) and Q-Q plots (for normality of residuals). Whenever residuals were not homogeneously distributed, appropriate heterogeneous variance models were fitted to meet the model assumptions by specifying a random/group statement (group = genotype *or* inoculation treatment *or* time point) after specifying the model statement. Bayesian information criterion (BIC) was used to determine the most suitable model that best fit data after accounting for model assumptions. Means separations were carried out using the PROC GLMMIX procedure of SAS.

## Results

### Differential gene expression analysis

DESeq2 analysis revealed 2317, 7133, and 432 differentially expressed genes (DEG) at 2, 7, and 30 DPI, respectively. However, only 588, 1718, and 100 of them had annotated metabolic pathways. The 588 and 1718 genes were constituents of 14 and 106 significantly enriched metabolic pathways while no significantly enriched pathway was observed at 30 DPI. As the greatest differential expression and pathway enrichment were observed at 7 DPI, we mainly focus on 7 DPI data for this paper. We identified 68 differentially expressed, lipid metabolism related genes at 7 DPI (Figure 1a, supplementary table 2).

**Figure 1.**
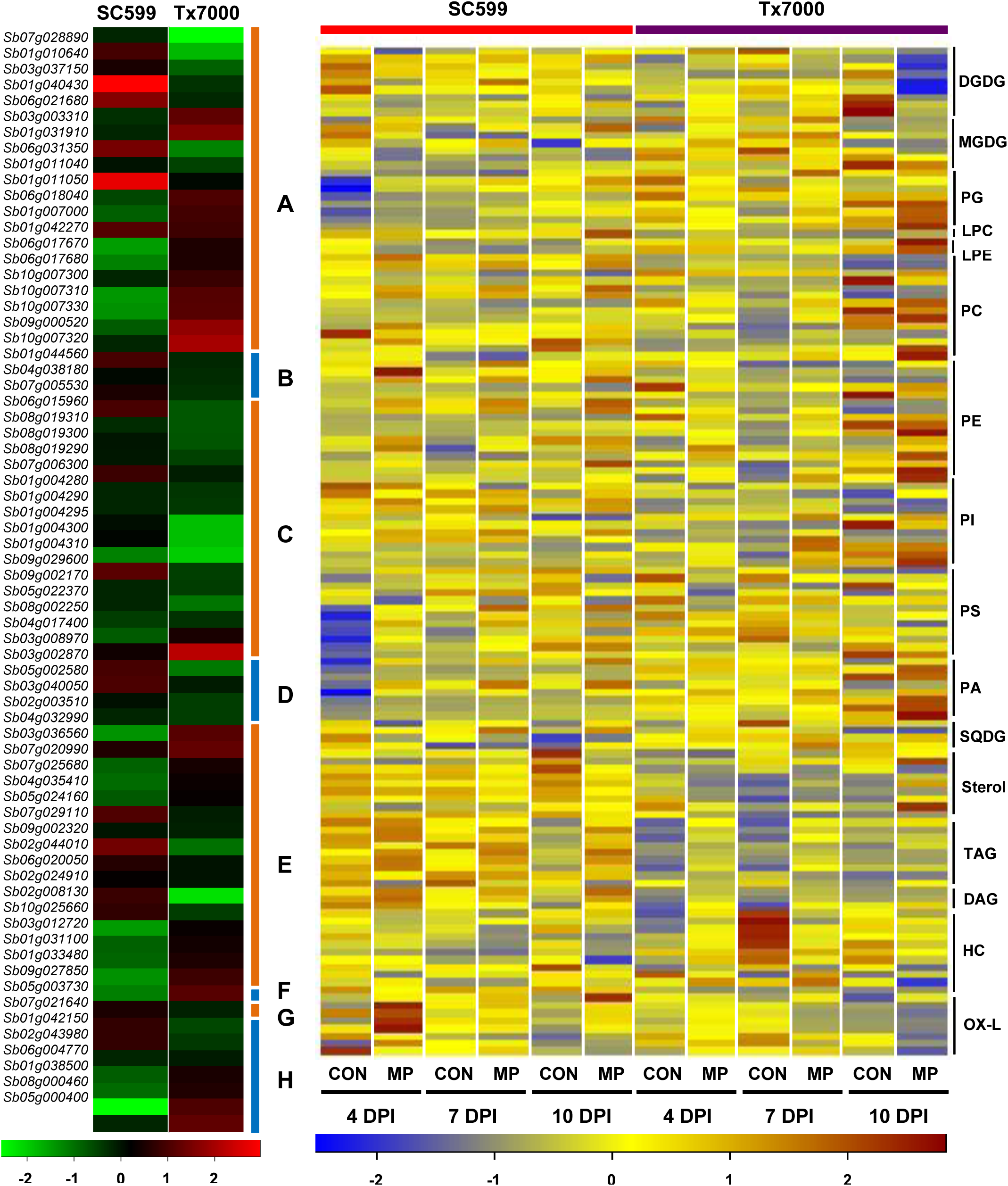
Comparative transcriptome and lipidome analysis of charcoal rot resistant and susceptible sorghum genotypes. (a) Transcriptome heat map depicting differentially expressed genes (log 2 fold) between charcoal rot resistant (SC599) and susceptible (Tx7000) sorghum genotypes in response to *Macrophomina phaseolina* at 7 days post-inoculation. Red, green, and black colors respectively represent up-regulated, down-regulated, and non-differentially expressed genes after *M. phaseolina* inoculation in comparison to mock-inoculated control treatment with sterile phosphate buffered saline. Each genotype contains three replicates per treatment. Letters from A to H respectively represent genes involved in jasmonic acid, trans, trans-farnesyl diphosphate, phytosterol, brassinosteroid, phosphatidic acid, and phosphatidylserine biosynthesis, MGDG to DGDG conversion, and phospholipid/glycolipid desaturation. (b) Lipidome heat map depicting mean concentrations (normalized intensity (%) per mg stalk dry weight) of 132 lipid species for SC599 and Tx7000 sorghum genotypes after receiving *M. phaseolina* (MP) and mock-inoculated control (CON) treatments at 4, 7, and 10 days post-inoculation (DPI). Each genotype contains five replicates per treatment. Blue, yellow, and maroon colors respectively represent low, medium, and high lipid concentrations. Abbreviations: DGDG, digalactosyldiacylglycerol; MGDG, monogalactosyldiacylglycerol; PG, phosphatidylglycerol; LPC, lyso-phosphatidylcholine; LPE, lyso-phosphatidylethanolamine; PC, phosphatidylcholine; PE, phosphatidylethanolamine; PI, phosphatidylinositol; PS, phosphatidylserines; PA, phosphatidic acid; SQDG, sulfoquinovosyldiacylglycerol; TAG, triacylglycerols; DAG, diacylglycerols; HC, hexosyl ceramides; OX-L, Ox-lipids.

Out of 68 DEGs, 20 were involved in jasmonic acid (JA) biosynthesis. Out of these 20, seven (*Sb01g010640*, *Sb01g031910*, *Sb01g040430, Sb03g003310*, *Sb03g037150*, *Sb06g021680*, and *Sb07g028890*) encoded for phospholipase A2. Except for *Sb03g003310* and *Sb01g031910*, other genes were significantly down-regulated in Tx7000 after pathogen inoculation with a 14.1 net log2 fold down-regulation. Although many of these genes were not significantly differentially expressed in SC599, *Sb01g040430* and *Sb06g021680* were significantly up-regulated (net log2 fold change = +4.0) upon pathogen inoculation. Out of four genes that encode lipoxygenase, *Sb01g011040* and *Sb06g031350* were down-regulated while *Sb06g018040* was up-regulated in Tx7000 after pathogen inoculation (net log2 fc = −4.4). *Sb01g011050* was significantly up-regulated (log2 fc = 2.5) in pathogen-inoculated SC599 while the other three genes were not significantly differentially expressed. Two cytochrome P450 74A3 genes (*Sb01g007000* and *Sb01g042270*; net log2 fc = +4.7) and seven 12-oxophytodienoate reductase genes (*Sb06g017670*, *Sb06g017680*, *Sb09g000520*, *Sb10g007300*, *Sb10g007310*, *Sb10g007320*, *Sb10g007330*; net log2 fc = +21.7) were significantly up-regulated in Tx7000 after pathogen inoculation while none of those were significantly differentially expressed in SC599.

Out of 68 DEGs, three were involved in the trans, trans-farnesyl diphosphate biosynthesis. Trans, trans-farnesyl diphosphate is the first precursor for phytosterol biosynthesis. Genes in this pathway (*Sb01g044560*, prenyltransferase; *Sb04g038180*, para-hydroxybenzoate-polyprenyl transferase; *Sb07g005530*, polyprenyl synthetase) were significantly down-regulated in Tx7000 upon *M. phaseolina* inoculation while none of those were significantly differentially expressed in SC599.

Sixteen genes out of 68 DEGs were related to phytosterol biosynthesis (campesterol, stigmasterol, and sitosterol). Five of these sixteen represented cycloartenol synthase (*Sb06g015960*, *Sb08g019310, Sb08g019300*, *Sb08g019290*, and *Sb07g006300*) and all of them were significantly down-regulated in Tx7000 after pathogen inoculation (net log2 fc = −11.5). Another six 24-methylenesterol C-methyltransferase 2 genes (*Sb01g004280*, *Sb01g004290*, *Sb01g004295*, *Sb01g004300*, *Sb01g004310*, *Sb09g029600*) were significantly down-regulated in pathogen inoculated Tx7000 (net log2 fc = −26.9). A cycloeucalenol cycloisomerase gene (*Sb09g002170*; log2 fc = −2.2) and two cytochrome P450 51 genes (*Sb05g022370*, *Sb08g002250*; net log2 fc = −5.9) were also significantly down-regulated in pathogen inoculated Tx7000. Two genes (*Sb04g017400*, log2 fc = +1.0; *Sb03g008970*, log2 fc = +6.6) that encode for C-14 sterol reductase (sterol delta-7 reductase) and 3-beta-hydroxysteroid-delta-isomerase, respectively were significantly up-regulated in *M. phaseolina*-inoculated Tx7000. None of the sixteen genes involved in phytosterol biosynthesis were significantly differentially expressed in SC599 due to pathogen inoculation.

Out of 68 DEGs, four were involved in the brassinosteroid biosynthesis. Two of those were steroid 22-alpha hydroxylase genes (*Sb03g002870* and *Sb05g002580*) and were significantly down-regulated in Tx7000 upon pathogen inoculation (net log2 fc = −5.2). The other two genes (*Sb03g040050* and *Sb02g003510*) encoded 3-oxo-5-alpha-steroid 4-dehydrogenase and were also found to be significantly down-regulated in pathogen-inoculated Tx7000 (net log2 fc = −4.3). None of these four genes were significantly differentially expressed in SC599 due to pathogen inoculation.

Seventeen out of 68 DEGs were responsible for phosphatidic acid biosynthesis. Seven of these encoded for diacylglycerol kinase (*Sb03g036560*, *Sb04g032990*, *Sb04g035410*, *Sb05g024160*, *Sb07g020990*, *Sb07g025680*, and *Sb07g029110*). The former five were significantly up-regulated in Tx7000 upon pathogen inoculation while the latter two were significantly down-regulated. The net log2 fold diacylglycerol kinase up-regulation was 6.8. None of these eight genes were significantly differentially expressed in SC599 in response to pathogen inoculation. Another three genes encoded phospholipase C (*Sb02g044010*, *Sb06g020050*, and *Sb09g002320*) and were significantly down-regulated in *M. phaseolina-*inoculated Tx7000 (net log2 fc = −5.3) while *Sb09g002320* was significantly up-regulated (log2 fc = +1.3) in SC599 after pathogen inoculation. Six genes represented phospholipase D (*Sb01g031100*, *Sb01g033480*, *Sb02g008130*, *Sb02g024910*, *Sb03g012720*, and *Sb10g025660*). Out of these, *Sb02g008130* and *Sb02g024910* were significantly down-regulated in Tx7000 after pathogen inoculation while the latter three were significantly up-regulated (net log2 fc = −5.6). Except for *Sb10g025660* (log2 fc = −1.0), the other five phospholipase D genes were not significantly differentially expressed in SC599 after pathogen inoculation.

A phosphatidylserine synthase gene (*Sb09g027850*) was significantly up-regulated (log2 fc = +3.1) in pathogen-inoculated Tx7000 while a digalactosyldiacylglycerol synthase gene (*Sb05g003730*) responsible for monogalactosyldiacylglycerol to digalactosyldiacylglycerol conversion was significantly down-regulated (log2 fc = −1.1). None of the two genes were significantly differentially expressed in SC599 upon pathogen inoculation.

Seven genes (*Sb01g038500*, *Sb01g042150*, *Sb02g043980, Sb05g000400*, *Sb06g004770*, *Sb07g021640*, and *Sb08g000460*) out of 68 DEGs were involved in phospholipid and glycolipid desaturation and encoded for Omega-6/-3 fatty acid desaturase. The former three genes were significantly down-regulated in pathogen-inoculated Tx7000 while the rest was significantly up-regulated. The net log2 fold up-regulation was 3.0. None of the seven genes were significantly differentially expressed in SC599 after pathogen inoculation.

### Lipidome analysis

The heat map shown in figure 1b depicts the mean concentrations (normalized intensity (%) per mg stalk dry weight) of 132 lipid species for SC599 and Tx7000 sorghum genotypes after receiving *M. phaseolina* (MP) and mock-inoculated control (CON) treatments at 4, 7, and 10 days post-inoculation (DPI). A profile analysis (Fig. 2) was conducted to examine the stalk lipid composition of two sorghum genotypes (SC599 and Tx7000) under control treatment across three time points. Although relative amounts differ between genotypes, in broader terms, phospholipids (∑ PG, PC, PE, PI, PS, PA) constituted the highest quantities (%) in both genotypes followed by hexosylceramide, galactolipids (∑ DGDG, MGDG, SQDG), phytosterols (∑ sterol glucosides, acyl(18:2) sterol glucosides, acyl(16:0) sterol glucosides), di/triacylglycerol (∑ NL297(18:2) containing TAG, NL295 (18:3) containing DAG/TAG, NL273 (16:0) containing DAG/TAG), lysophospholipids (∑ LysoPC, LysoPE), and Ox-lipids (∑ prec291 (18:3-2O) or 18:4-O, prec293 (18:2-2O) or 18:3-O) respectively.

**Figure 2.**
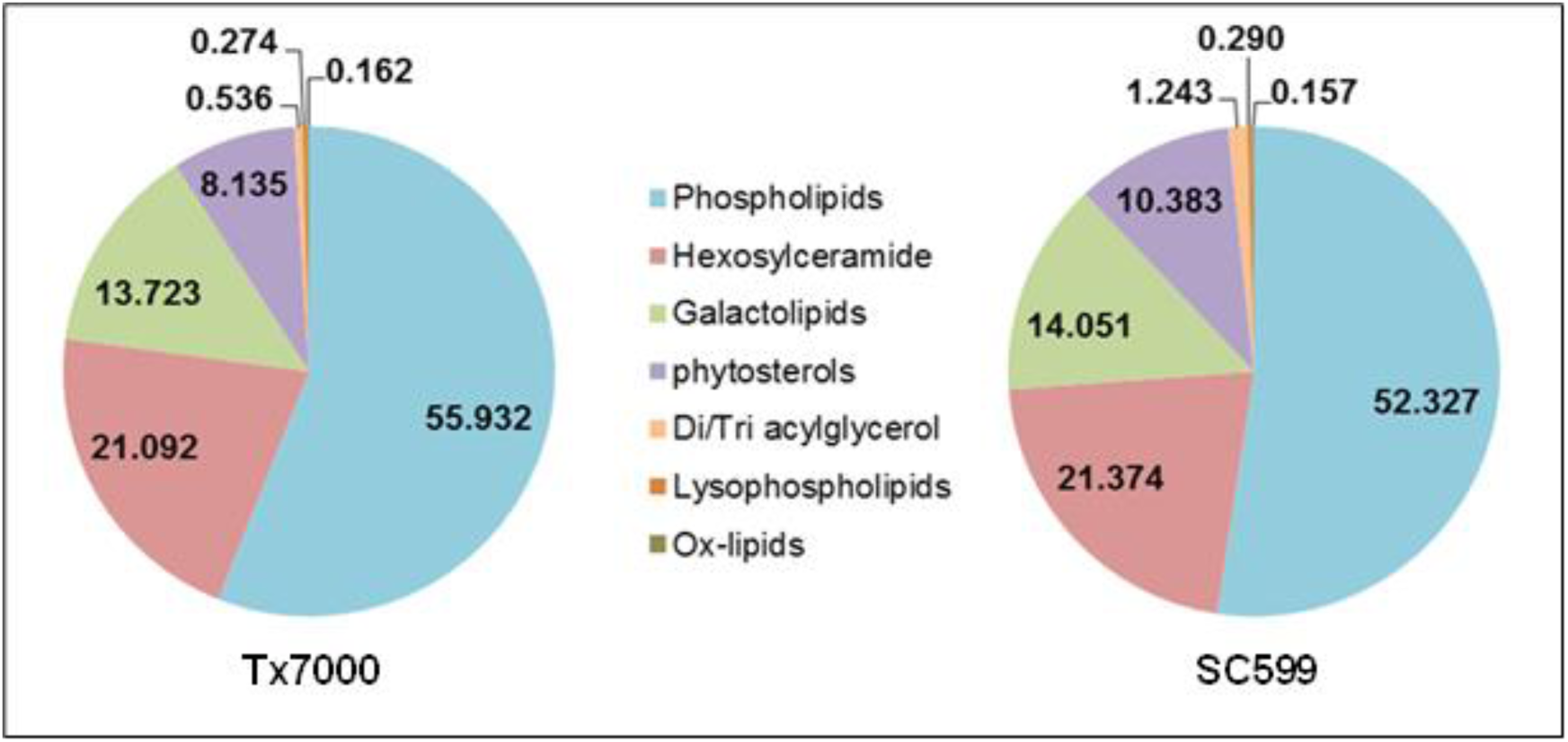
Stalk lipid composition (%) of two tested sorghum genotypes (SC599 and Tx7000) after control treatment across three time points (4, 7, 10 days post-inoculation). Phospholipids = ∑ (PG, PC, PE, PI, PS, PA); galactolipids = ∑ (DGDG, MGDG, SQDG); phytosterols = ∑ (sterol glucosides, acyl(18:2) sterol glucosides, acyl(16:0) sterol glucosides); Di/Triacylglycerol = ∑ (NL297(18:2) containing TAG, NL295 (18:3) containing DAG/TAG, NL273 (16:0) containing DAG/TAG); lysophospholipids = ∑ (LysoPC, LysoPE), and Ox-lipids = ∑ (prec291 (18:3-2O) or 18:4-O, prec293 (18:2-2O) or 18:3-O)). ∑ = sum.

### Analysis of lipid classes

The genotype × inoculation treatment interaction effect was significant across three time points for MGD:DGDG ratio, PS, sterol glucosides (∑ campesterol-glc, stigmasterol-glc, sitosterol-glc), acyl (18:2) sterol glucosides (∑ campesterol-glc(18:2), stigmasterol-glc(18:2), sitosterol-glc(18:2)), and total ox-lipids (∑ PE(16:0/18:3-2O), MGDG(18:4-O/18:3), PC(16:0/18:3-2O), MGDG(18:3-2O/18:3), PE(18:2/18:3-O), PE(18:2/18:2-2O), and PC(16:0/18:3-O)) (Table 1). Compared to control treatment, pathogen inoculation significantly increased the MGDG:DGDG ratio of Tx7000 (*P* = 0.0004) but did not affect this ratio in SC599 (*P* = 0.7288) (Fig. 3a). *M. phaseolina* significantly decreased the PS content of Tx7000 (*P* = 0.0136) but did not affect the PS content in SC599 (*P* = 0.3882) (Fig. 3b). *M. phaseolina* also significantly reduced the sterol glucosides content of Tx7000 (*P* = 0.0031) while no significant impact was observed in SC599 (*P* = 0.7141) (Fig. 3c). Although pathogen inoculation did not significantly affect acyl (18:2) sterol glucoside content in Tx7000 (*P* = 0.9184), a significant reduction was observed in SC599 (*P* = 0.0012) (Fig. 3d). *M. phaseolina* inoculation significantly increased the total ox-lipid content in SC599 (*P* = 0.0148) while it significantly decreased ox-lipid in Tx7000 (*P* = 0.0309) (Fig. 3e).

**Figure 3.**
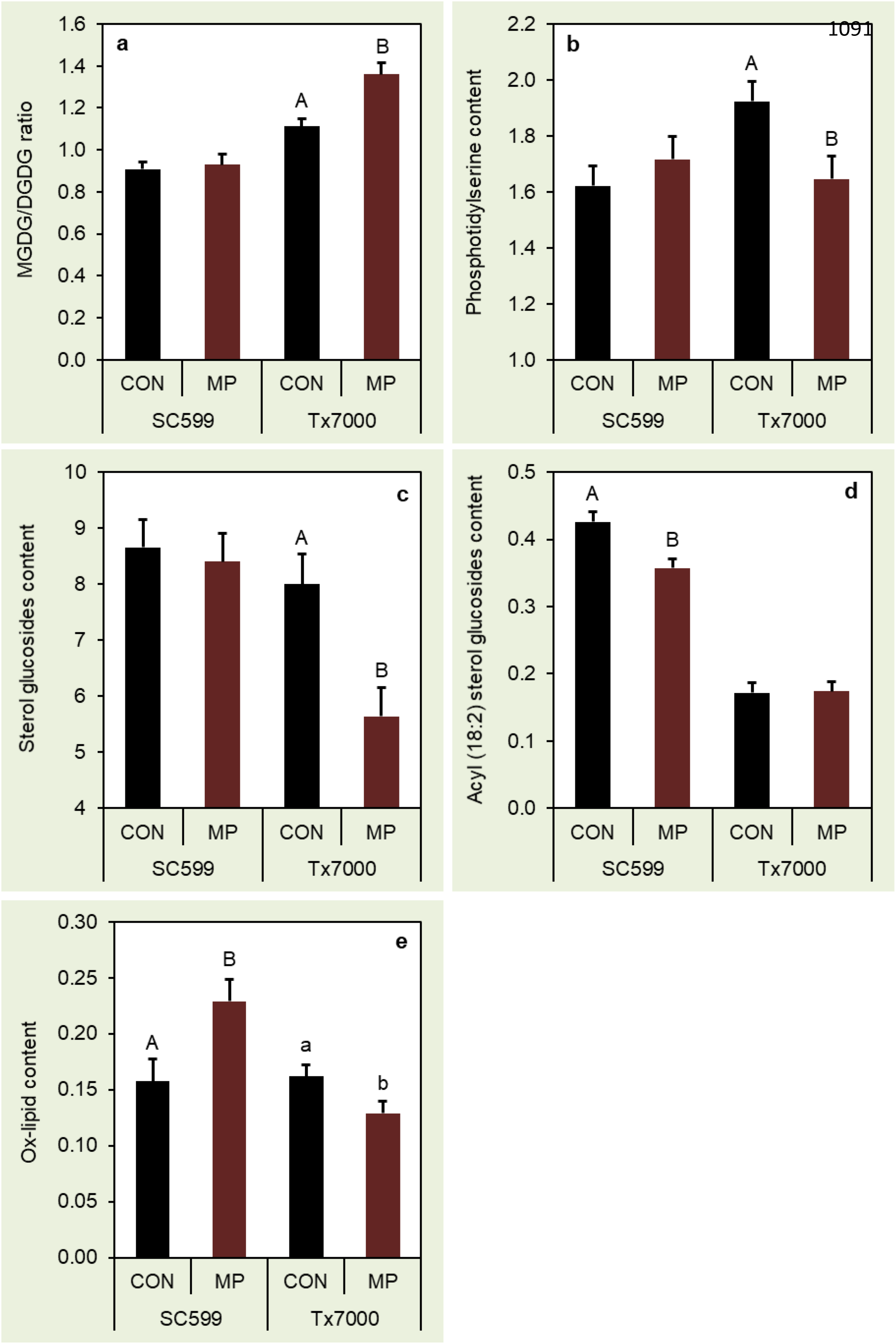
Comparison of the mean values (normalized mass spectral signal per mg of stalk tissue) among inoculation treatments for (a) monogalactosyldiacylglycerol(MGDG)/ digalactosyldiacylglycerol (DGDG) ratio, (b) phosphatidylserine, (c) sterol glucoside, (d) acyl(18:2) sterol glucoside, and (e) ox-lipid content at each genotype across three time points (4, 7, and 10 days post-inoculation). Means followed by different letters within each genotype are significantly different while the treatments without letter designations within each genotype are not significantly different at α = 0.05. Error bars represent standard errors. CON = phosphate-buffered saline mock-inoculated control, MP = *Macrophomina phaseolina*.

**Table 1.** F-statistic *P*-values from analysis of variance (ANOVA) for different sorghum stalk lipid classes isolated from charcoal-rot-resistant (SC599) and susceptible (Tx7000) sorghum genotypes after inoculation with *M. phaseolina* and phosphate buffered saline (mock-inoculated control) at three post-inoculation time points (4, 7, and 10 days post-inoculation) (α = 0.05). Lipids were analyzed using an electrospray ionization-triple quadrupole mass spectrometer.

| Lipid class / effect | *P*-value |  |  |  |  |  |  |
| --- | --- | --- | --- | --- | --- | --- | --- |
|  | Genotype (G) | Time (T) | G × T | Trt* (I) | G × I | T × I | G × T × I |
| DGDG† | 0.0002 | 0.1330 | 0.3049 | 0.0302 | 0.6788 | 0.2693 | 0.0275 |
| MGDG | 0.1025 | 0.0048 | 0.6768 | 0.9439 | 0.9147 | 0.4966 | 0.0638 |
| MGDG/DGDG | <0.0001 | 0.0106 | 0.2427 | 0.0042 | **0.0143** | 0.5874 | 0.9198 |
| SQDG | 0.8342 | 0.1793 | 0.8570 | 0.4282 | 0.8711 | 0.2635 | 0.0213 |
| PG | <0.0001 | 0.0278 | 0.0022 | 0.8903 | 0.0284 | 0.9771 | 0.0214 |
| PC | 0.8012 | 0.1981 | 0.5463 | 0.0695 | 0.6494 | 0.2462 | 0.2508 |
| PE | 0.0266 | 0.2560 | 0.1483 | 0.2908 | 0.4520 | 0.4450 | 0.2195 |
| PE/PC | 0.0005 | 0.4360 | 0.0482 | 0.8763 | 0.0735 | 0.6460 | 0.3786 |
| PI | 0.0110 | 0.0085 | 0.3554 | 0.0204 | 0.3264 | 0.3787 | 0.2057 |
| PS | 0.1392 | 0.0118 | 0.1097 | 0.2378 | **0.0190** | 0.2142 | 0.0765 |
| PA | <0.0001 | 0.0211 | 0.2362 | 0.0002 | 0.9484 | 0.5027 | 0.0243 |
| Galactolipids°/Phospholipids‡ | 0.0288 | 0.0081 | 0.1794 | 0.0142 | 0.9596 | 0.8798 | 0.0009 |
| LysoPC | <0.0001 | 0.0521 | 0.6556 | 0.1929 | 0.4753 | 0.0848 | 0.3285 |
| LysoPE | 0.0936 | 0.1601 | 0.5967 | 0.2781 | 0.5530 | 0.3353 | 0.9535 |
| Sterol Glucosides | 0.0019 | 0.5439 | 0.7711 | 0.0145 | **0.0464** | 0.9666 | 0.8195 |
| Acyl(18:2)Sterol Glucosides | <0.0001 | 0.0811 | 0.5158 | 0.0218 | **0.0152** | 0.6307 | 0.0624 |
| Acyl(16:0)Sterol Glucosides | <0.0001 | 0.1510 | 0.0657 | 0.3174 | 0.3141 | 0.1602 | 0.0803 |
| NL297(18:2)containing TAG | <0.0001 | 0.3251 | 0.0473 | 0.0023 | 0.7659 | 0.9300 | 0.1361 |
| NL295(18:3)containing DAG,TAG | <0.0001 | 0.4359 | 0.3500 | <0.0001 | 0.1266 | 0.9599 | 0.1738 |
| NL273(16:0)containing DAG,TAG | <0.0001 | 0.1934 | 0.0976 | 0.0004 | 0.5034 | 0.7124 | 0.1367 |
| HexCer | 0.7335 | 0.7213 | 0.3913 | 0.0464 | 0.9281 | 0.2743 | 0.4609 |
| Prec291(18:3-2O)or18:4-O (A) | <0.0001 | 0.0026 | 0.5462 | 0.0128 | **0.0013** | 0.0136 | 0.7611 |
| Prec293(18:2-2O)or18:3-O (B) | 0.9411 | <0.0001 | 0.0151 | 0.1397 | **0.0165** | 0.6564 | 0.2161 |
| Total ox-lipids (A+B) | 0.0043 | <0.0001 | 0.1882 | 0.2308 | **0.0020** | 0.0585 | 0.9105 |

\*Trt = Inoculation treatment. † DGDG = digalactosyldiacylglycerol; MGDG = Monogalactosyldiacylglycerol; SQDG = Sulfoquinovosyl diacylglycerol; PG = Phosphatidylglycerol; PC = Phosphatidylcholine; PE = Phosphatidylethanolamine; PI = Phosphatidylinositol; PS = Phosphatidylserine; PA = Phosphatidic acid; LysoPC = Lysophosphatidylcholine; LysoPE = Lysophosphatidylethanolamine; TAG = Triacylglycerol; DAG = Diacylglycerol; HexCer = Hexosylceramide. °Galactolipids = (DGDG + MGDG + SQDG). ‡Phospholipids = (PG + PC + PE + PI + PS + PA).

PA analysis revealed significantly increased PA content in SC599 after *M. phaseolina* inoculation at 4 DPI (*P* < 0.0001). MP did not significantly affect PA content in Tx7000 (*P* = 0.9166) (Fig. 4ai). The main effects of genotype and inoculation treatment on PA were evident across 7 and 10 DPI. Tx7000 had significantly greater PA content than SC599 (*P* < 0.0001) across the two inoculation treatments, and across 7 and 10 DPI (Fig. 4aii). Compared to control, *M. phaseolina* significantly increased PA content across two genotypes, and across 7 and 10 DPI (*P* = 0.0007) (Fig. 4aiii). PG content was significantly greater in SC599 after *M. phaseolina* inoculation (*P* = 0.047) at 4 DPI while the same was significantly lower in Tx7000 after *M. phaseolina* inoculation (*P* = 0.0173) (Fig. 4bi). The main effect of genotype for PG was evident across inoculation treatments and, across 7 and 10 DPI where Tx7000 had a significantly greater PG content than SC599 (*P* = 0.0001) (Fig. 4bii). The galactolipids:phospholipids ratio was significantly decreased in SC599 after MP inoculation at 4 DPI (*P* = 0.0125) while it significantly increased the ratio in Tx7000 (*P* = 0.0491) (Fig. 4ci). Although *M. phaseolina* inoculation did not significantly affect the galactolipids:phospholipids ratio of SC599 (*P* = 0.8377) across 7 and 10 DPI, it significantly decreased the galactolipids:phospholipids ratio in Tx7000 (*P* = 0.0074) (Fig. 4cii).

**Figure 4.**
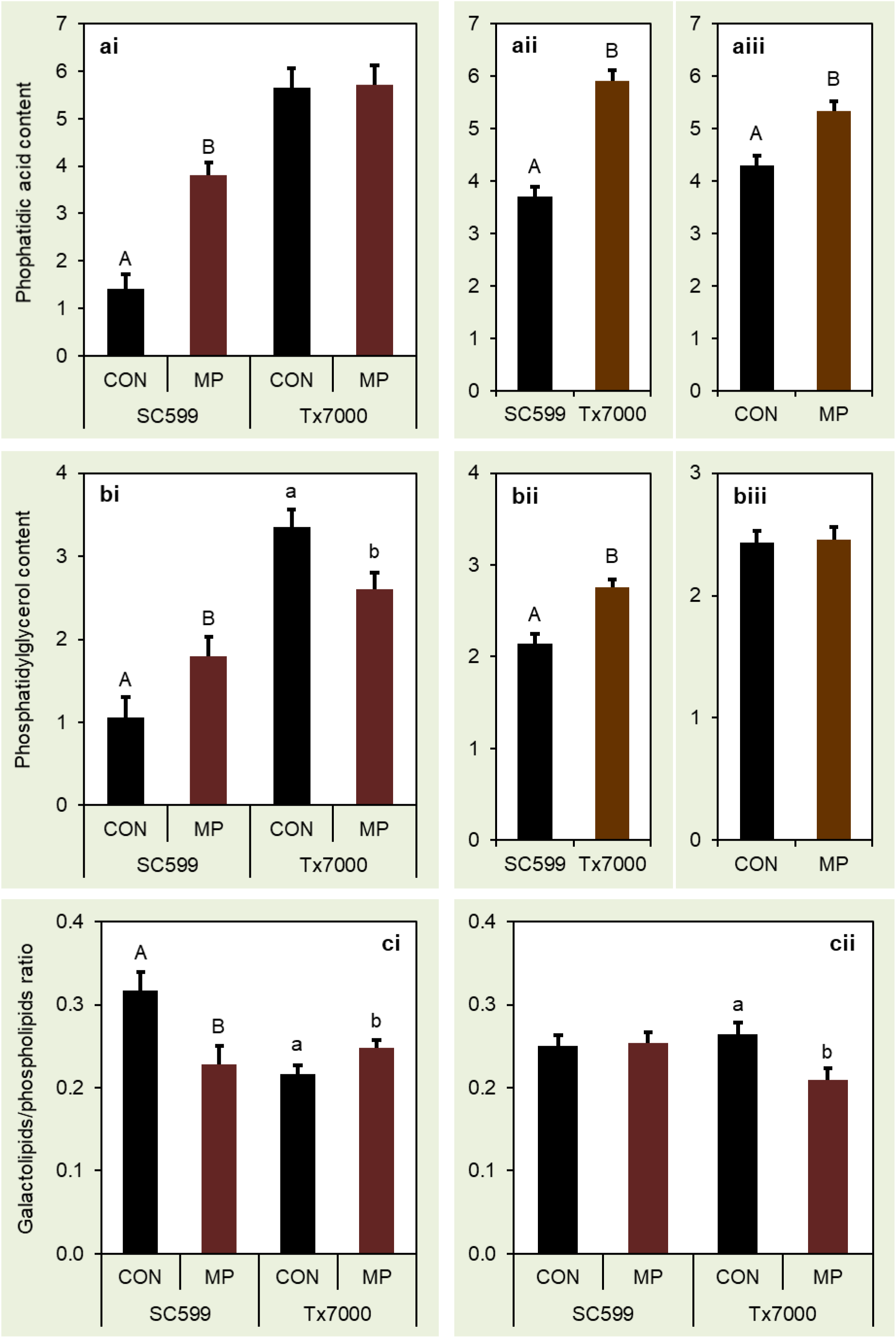
Comparison of the mean (normalized mass spectral signal per mg of stalk tissue) phosphatidic acid content (ai) at 4 days post-inoculation (DPI) (aii) across 7 and 10 DPI and inoculation treatments (aiii) across 7 and 10 DPI and two genotypes; phosphatidylglycerol content (bi) at 4 DPI (bii) across 7 and 10 DPI and inoculation treatments (biii) across 7 and 10 DPI and two genotypes; and the galactolipids/phospholipid ratio (ci) at 4 DPI and (cii) across 7 and 10 DPI. Means followed by different letters (within each letter case) are significantly different while the treatments without letter designations are not significantly different at α = 0.05. Error bars represent standard errors. CON = phosphate-buffered saline mock-inoculated control, MP = *Macrophomina phaseolina*.

### Analysis of lipid species

Out of 132 lipid species analyzed, 31 showed significant genotype × inoculation treatment interaction across the three post-inoculation stages (Table 2). Compared to control, *M. phaseolina* inoculation significantly increased the PG(34:3) content of SC599 while it significantly decreased the same in Tx7000. *M. phaseolina* significantly increased the PG (36:2) in Tx7000. *M. phaseolina* inoculation significantly increased PC(34:3), PC(36:6), and PC(36:1) content of SC599 while it significantly decreased PC(34:2). A significant PC(36:6) and PC(36:2) increase was also observed in Tx7000 after *M. phaseolina* inoculation. *M. phaseolina* inoculation significantly increased the PE(34:4), PE(34:3), and PE(36:6) species in SC599. These species were not significantly affected in Tx7000 after pathogen inoculation. However, PE(36:2) and PE(42:2) species were present in significantly greater quantities in *M. phaseolina*-inoculated Tx7000 although *M. phaseolina* did not significantly affect PE(36:2) and PE(42:2) of SC599. *M. phaseolina* inoculated Tx7000 had significantly greater amounts of PI(34:2), PI(36:4), and PI(36:2) species. *M. phaseolina* significantly reduced the PI(34:2) in SC599 while did not significantly affect the PI(36:4) and PI(36:2). *M. phaseolina* inoculation significantly increased the PS(36:3), PS(38:3), PS(40:3) species in SC599 although none of them were significantly affected in Tx7000. PS(34:3) was significantly reduced in *M. phaseolina-*inoculated Tx7000. However, PS(34:3) was not significantly affected by *M. phaseolina* in SC599. Although *M. phaseolina* inoculation significantly increased the PA(34:3) and PA(36:6) species in both sorghum genotypes, there was a greater increase with SC599 compared to Tx7000. *M. phaseolina* significantly decreased campesterol-glc(18:2) in SC599 but not in Tx7000. The stigmasterol-glc:sitosterol-glc ratio was significantly greater in pathogen-inoculated Tx7000. However, pathogen inoculation did not significantly affect the ratio of SC599. Although *M. phaseolina* significantly increased TAG(18:3/36:9) and TAG(16:0/36:6) in both sorghum genotypes, there was a greater increase in SC599. SC599 had significantly lower DAG(34:2) content after *M. phaseolina* inoculation although the same was not significantly affected in Tx7000. The ox-lipid species MGDG(18:4-O/18:3), MGDG(18:3-2O/18:3), PE(16:0/18:3-2O), PE(18:2/18:2-2O), and PC(16:0/18:3-2O) were found to be significantly greater in SC599 after *M. phaseolina* inoculation. Except for the significantly reduced PE(18:2/18:2-2O), *M. phaseolina* did not affected the aforementioned ox-lipid species in Tx7000.

**Table 2.** Lipid species with significant genotype × inoculation treatment interactions across three post-inoculation stages (4, 7, and 10 days post-inoculation). Mean lipid content (normalized signal in % basis) and *P*-values for mean difference between control and *Macrophomina phaseolina* are given (α = 0.05).

| Lipid species | P-value<br>$G \times I^*$ | SC599 | | | Tx7000 | | |
| --- | --- | --- | --- | --- | --- | --- | --- |
|  |  | Mean |  | P-value | Mean |  | P-value |
|  |  | CON<sup>‡</sup> | MP<sup>•</sup> |  | CON | MP |  |
| PG(34:3)<sup>†</sup> | 0.0006 | 0.519 | 0.654 | 0.0127 | 0.699 | 0.626 | 0.0015 |
| PG(36:2) | 0.0038 | 0.007 | 0.008 | 0.7542 | 0.014 | 0.024 | 0.0005 |
| PC(34:3) | 0.0003 | 6.164 | 7.540 | <0.0001 | 4.074 | 4.358 | 0.1321 |
| PC(34:2) | 0.0127 | 9.554 | 8.667 | 0.0178 | 10.544 | 11.010 | 0.2234 |
| PC(36:6) | 0.0001 | 0.899 | 1.405 | <.0001 | 0.434 | 0.531 | 0.0288 |
| PC(36:2) | 0.0026 | 0.392 | 0.413 | 0.4497 | 0.516 | 0.733 | 0.0003 |
| PC(36:1) | 0.0115 | 0.058 | 0.081 | 0.0047 | 0.063 | 0.054 | 0.3268 |
| PE(34:4) | 0.0485 | 0.010 | 0.014 | 0.0018 | 0.007 | 0.008 | 0.0905 |
| PE(34:3) | 0.0013 | 1.953 | 2.553 | <.0001 | 1.548 | 1.597 | 0.6338 |
| PE(36:6) | 0.0006 | 0.181 | 0.319 | <.0001 | 0.105 | 0.127 | 0.0869 |
| PE(36:2) | 0.0002 | 0.072 | 0.077 | 0.2534 | 0.101 | 0.153 | <.0001 |
| PE(42:2) | 0.0380 | 0.131 | 0.140 | 0.1744 | 0.151 | 0.182 | 0.0003 |
| PI(34:2) | 0.0130 | 4.591 | 4.154 | 0.0435 | 4.848 | 5.399 | 0.0347 |
| PI(36:4) | 0.0290 | 0.178 | 0.162 | 0.4206 | 0.230 | 0.278 | 0.0218 |
| PI(36:2) | <0.0001 | 0.061 | 0.065 | 0.2765 | 0.080 | 0.144 | <.0001 |
| PS(34:3) | 0.0190 | 0.330 | 0.356 | 0.2664 | 0.248 | 0.198 | 0.0206 |
| PS(36:3) | 0.0010 | 0.036 | 0.047 | <.0001 | 0.035 | 0.032 | 0.3687 |
| PS(38:3) | 0.0020 | 0.061 | 0.080 | 0.0004 | 0.066 | 0.060 | 0.2986 |
| PS(40:3) | 0.0095 | 0.102 | 0.126 | 0.0172 | 0.134 | 0.115 | 0.1361 |
| PA(34:3) | 0.0041 | 0.700 | 1.122 | <.0001 | 0.817 | 0.981 | <.0001 |
| PA(36:6) | 0.0002 | 0.070 | 0.144 | <.0001 | 0.060 | 0.080 | 0.0002 |
| Campesterol-Glc(18:2) | 0.0276 | 0.155 | 0.132 | 0.0078 | 0.066 | 0.069 | 0.7538 |
| Stigmasterol-Glc/Sitosterol-Glc | 0.0162 | 0.393 | 0.392 | 0.9907 | 0.559 | 0.830 | 0.0062 |
| TAG(18:3/36:9) | 0.0006 | 0.076 | 0.190 | <.0001 | 0.016 | 0.037 | 0.0050 |
| TAG(16:0/36:6) | 0.0016 | 0.067 | 0.140 | <.0001 | 0.016 | 0.032 | 0.0051 |
| DAG(34:2) | 0.0026 | 0.021 | 0.015 | <.0001 | 0.017 | 0.017 | 0.6329 |
| MGDG(18:4-O/18:3) | 0.0188 | 0.016 | 0.025 | 0.0302 | 0.009 | 0.007 | 0.3356 |
| MGDG(18:3-2O/18:3) | 0.0056 | 0.014 | 0.025 | 0.0063 | 0.011 | 0.009 | 0.4408 |
| PE(16:0/18:3-2O) | 0.0008 | 0.009 | 0.026 | 0.0014 | 0.009 | 0.006 | 0.2419 |
| PE(18:2/18:2-2O) | 0.0001 | 0.021 | 0.030 | 0.0144 | 0.036 | 0.026 | 0.0011 |
| PC(16:0/18:3-O) | 0.0418 | 0.013 | 0.010 | 0.0321 | 0.008 | 0.010 | 0.4566 |
| PC(16:0/18:3-2O) | 0.0006 | 0.008 | 0.021 | 0.0003 | 0.007 | 0.006 | 0.6307 |

\* $G \times I$ = Genotype by Inoculation treatment interaction. <sup>‡</sup>CON = Mock inoculated control treatment. <sup>•</sup>MP = *Macrophomina phaseolina* treatment. <sup>†</sup>PG = Phosphatidylglycerol; PC = Phosphatidylcholine; PE = Phosphatidylethanolamine; PI = Phosphatidylinositol; PS = Phosphatidylserine; PA = Phosphatidic acid; TAG = Triacylglycerol; DAG = Diacylglycerol; MGDG = Monogalactosyldiacylglycerol

## Discussion

### Increased MGDD/DGDG ratio and charcoal rot susceptibility

The enzyme digalactosyldiacylglycerol (DGD) synthase transfers a galactose from UDP-galactose onto MGDG to form DGDG (Boudière *et al*., 2014). MGDG does not form bilayers in mixtures with water while DGDG is a bilayer-forming lipid (Webb & Green, 1991). The ratio of non-bilayer-forming to bilayer-forming lipids is critical for protein folding and insertion (Gounaris & Barber, 1983; Bogdanov & Dowhan, 1999) as well as for intracellular protein trafficking (Kusters *et al*., 1994). Therefore, the MGDG:DGDG ratio in chloroplasts must be tightly regulated to maintain a stable physical phase and for the proper functioning of the thylakoid membranes. The decreased ratio of MGDG:DGDG enhances the stability of the membrane under various stresses (Moellering *et al*, 2010; Chen *et al*., 2006; Hazei & Williams, 1990; Welti *et al*., 2002). In the current study, DGD synthase gene was significantly down-regulated in Tx7000 after *M. phaseolina* inoculation which contributed to impeded MGDG to DGDG conversion. This was evident with the increased MGDD:DGDG ratio observed in Tx7000 upon *M. phaseolina* inoculation. Therefore, *M. phaseolina* may promote charcoal rot susceptibility through a negative impact on thylakoid membrane stability and function.

### Decreased phosphatidylserine (PS) content and charcoal rot susceptibility

The two most abundant classes of phospholipids in plant mitochondria and cell membrane are phosphatidylcholine (PC) and phosphatidylethanolamine (PE) (Horvath & Daum, 2013). Maintaining a lower PE:PC ratio is important to enhance the stability of membranes under various stresses (Moellering *et al*., 2010; Chen *et al*., 2006; Hazei & Williams, 1990; Welti *et al*., 2002). In the current study, we did not observe a significant alteration in this ratio after pathogen inoculation in either genotype. However, phosphatidylserine (PS) is also considered an important lipid constituent in cell and mitochondria membranes (Horvath & Daum, 2013). Although a relatively minor plant cell lipid class (Devaiah *et al*., 2006; Nakamura & Ohta, 2007), PS plays an important role in cell death signaling, vesicular trafficking, lipid–protein interactions, and membrane lipid metabolism (Vance, 2008). We observed a significant reduction in PS in Tx7000 as a lipid class and three PS species (PS(36:3), PS(38:3), and PS(40:3)) increment in SC599 after *M. phaseolina* inoculation. The significantly up-regulated phosphatidylserine synthase gene in Tx7000 after *M. phaseolina* inoculation may be an indication of Tx7000’s need to produce extra PS. Therefore, despite the unaltered PE:PC ratio, the current study provided some evidence for the potential negative impacts of *M. phaseolina* on mitochondrial and cell membrane stability and functioning in the charcoal-rot-susceptible genotype, Tx7000.

### MP decrease phytosterol biosynthesis in Tx7000 while increasing its stigmasterol/sitosterol ratio

Phytosterols (campesterol, stigmasterol, and sitosterol) represent the most abundant sterols in plants (Benveniste, 2004). They are integral constituents of membrane lipid bilayer and regulators of membrane fluidity and permeability, and influence membrane properties, functions, and structure (Demel & De Kruyff, 1976; Bloch, 1983; Schuler *et al*., 1991; Schaller, 2003; Roche *et al*., 2008). Phytosterols also play a crucial role in plant innate immunity against phytopathogens. For instance, silencing of *N. benthamiana* squalene synthase, a key gene in phytosterol biosynthesis, compromised non-host resistance to a few pathovars of *Pseudomonas syringae* and *Xanthomonas campestris* while an *Arabidopsis* sterol methyltransferase mutant (sterol methyltransferase2) involved in sterol biosynthesis also compromised plant innate immunity against bacterial pathogens (Wang *et al*., 2012). In the current study, we observed significant down-regulation of prenyltransferase, para-hydroxybenzoate-polyprenyl transferase, and polyprenyl synthetase genes in Tx7000 after MP inoculation, which may have limited trans, trans-farnesyl diphosphate biosynthesis. Trans, trans-farnesyl diphosphate is essentially the primary precursor for the biosynthesis of all phytosterols (http://pathway.gramene.org/ gramene/sorghumcyc.shtml). Moreover, important genes involved in phytosterol biosynthesis such as cycloartenol synthase, 24-methylenesterol C-methyltransferase 2, cycloeucalenol cycloisomerase, and cytochrome P450 51 were significantly down-regulated in Tx7000 after *M. phaseolina* inoculation. Confirming the gene expression data, significantly lower sterol glucoside (∑ campesterol, stigmasterol, sitosterol) content was observed in Tx7000 after *M. phaseolina* inoculation. Therefore, *M. phaseolina* inoculation associated phytosterol reduction in Tx7000 could result in cell membrane destabilization and compromised plant immunity that could contribute to enhanced charcoal rot susceptibility.

The stigmasterol to sitosterol ratio was found to be significantly greater in *M. phaseolina* inoculated Tx7000. The sitosterol to stigmasterol conversion is triggered through perception of pathogen-associated molecular patterns such as flagellin and lipopolysaccharides, and the generation of reactive oxygen species (ROS) (Griebel & Zeier, 2010). Therefore, the increased stigmasterol to sitosterol ratio observed in the current study provided a clue about the strong oxidative stress experienced by Tx7000 after *M. phaseolina* inoculation. Previous studies provided evidence for *M. phaseolina*’s ability to create oxidative stress responses in charcoal-rot-susceptible sorghum genotypes such as Tx7000 and BTx3042 through induced host nitric oxide (NO) and ROS biosynthesis (this data has been submitted for publication and currently under review). Moreover, through mutant analysis and exogenous sterol application, Griebel & Zeier (2010) have shown that an increased stigmasterol to sitosterol ratio in *Arabidopsis* leaves weakens specific plant defence responses, which results in enhanced susceptibility against *Pseudomonas syringae*. Therefore, it is possible that *M. phaseolina* inoculation associated stigmasterol to sitosterol ratio increase could contribute to enhanced charcoal rot disease susceptibility in Tx7000.

### Brassinosteroid and charcoal rot disease reaction

The biosynthetic pathway for phytosterols also provides precursors for brassinosteroids, phytohormones involved in the regulation of plant growth and development (Fujioka *et al*., 2002). In the current study, genes involved in brassinosteroid biosynthesis such as steroid 22-alpha hydroxylase and 3-oxo-5-alpha-steroid 4-dehydrogenase were significantly down-regulated in Tx7000 after *M. phaseolina* inoculation. Therefore, although host brassinosteroid content was not directly measured in this study, the significantly lower sterol glucosides (precursors for brassinosteroids biosynthesis) content as well as the down-regulation of key genes involved in brassinosteroid biosynthesis suggested the bottleneck that could be faced by Tx7000 in synthesizing brassinosteroid after *M. phaseolina* inoculation. Perhaps, impeded brassinosteroid biosynthesis could be a mechanism through which Tx7000 attenuates further upsurge of ROS mediated oxidative stress after *M. phaseolina* inoculation. Brassinosteroids are reported to induce ROS accumulation and programmed cell death in plants (Fukuda, 2000; Kuriyama *et al*., 2001; Roberts *et al*., 2000; Xia *et al*., 2009).

### Ox-lipids, jasmonic acid, and charcoal rot disease reaction

Like oxylipins, ox-lipids may also function as signaling molecules that initiate stress responses in plants (Andersson *et al*., 2006). Plants can produce ox-lipids in response to a variety of stresses including pathogen attacks (Thoma *et al*., 2003). Ox-lipids are produced enzymatically through the action of lipoxygenase or non-enzymatically through the action of reactive oxygen species (ROS) (Zoeller *et al*., 2012). The primary product of lipoxygenase mediated lipid oxidation is lipid hydroperoxides while phytoprostanes are the primary product of ROS-mediated oxidation (Christensen & Kolomiets, 2011). The precursor lipid hydroperoxides can further be subjected to various enzymatic reactions which results in the generation of a variety of oxylipins including 12-oxo-phytodienoic acid (OPDA) and jasmonic acid (JA) (Imbusch & Mueller, 2000; Gobel *et al*., 2002; Mosblech *et al*., 2009). In the current study, compared to control, we observed significantly higher ox-lipid content in the charcoal-rot-resistant genotype, SC599, after *M. phaseolina* inoculation. Although not directly measured, the presence of ox-lipids in higher quantities (particularly PC(16:0/18:3-O) and PC(16:0/18:3-2O) species) along with the net up-regulation of phospholipase A2 and lipoxygenase genes (phospholipase A2 and lipoxygenase are necessary to produce lipid hydroperoxides which are essential precursors for JA biosynthesis) suggested SC599’s potential to produce ample amounts of jasmonic acid under *M. phaseolina* inoculation. JA is an important plant hormone which confers resistance against necrotrophic pathogens (McDowell & Dangl, 2000; Glazebrook, 2005). On the other hand, although genes involve in the latter steps of JA biosynthesis such as cytochrome P450 74A3 and 12-oxophytodienoate reductase are highly up-regulated in Tx7000 after *M. phaseolina* inoculation, the net down-regulation of the phospholipase A2 and lipoxygenase genes (needed for initial steps in JA biosynthesis) may hinder JA synthesis in Tx7000 after *M. phaseolina* infection. The impeded potential of Tx7000 to produce JA under *M. phaseolina* infection is also supported by its significantly lower ox-lipid content. As described earlier, Tx7000 appeared to experience a strong oxidative stress after *M. phaseolina* infection. Oxidative stress results in the biosynthesis of phytoprostanes from the available polyunsaturated fatty acid (PUFA, particularly linolenate), which reduces available linolenate pools to be utilized in the JA production. Therefore, Tx7000 appeared to suffer with the major precursor (linolenate) shortage to produce JA under *M. phaseolina* infection, which in turn could make Tx7000 more susceptible to this important necrotrophic fungus.

### Implications of phosphatidic acid (PA) in charcoal rot disease reaction

The phosphatidic acid (PA) involved in cell signaling is produced via two phospholipase pathways. It can be generated directly through the hydrolysis of structural phospholipids through phospholipase D (PLD) activity (Testerink & Munnik, 2005; Munnik, 2001; Wang, 2004). It is also synthesized via the successive action of phospholipase C (PLC) and diacylglycerol kinase (DAGK) (Testerink & Munnik, 2005; Munnik, 2001; Wang, 2004). In this pathway, PLC hydrolyzes phosphatidylinositol (PI) into inositol-1,4,5-trisphosphate and diacylglycerol (DAG). The resulting DAG is rapidly phosphorylated to PA by DAGK (Testerink & Munnik, 2005). In the current study, both genotypes contained significantly higher PA(34:3) and PA(36:6) upon *M. phaseolina* inoculation. However, compared to control, the resistant genotype had increased PA(34:3) and PA(36:6) after pathogen inoculation. The PLC/DAGK pathway contributes to this increase. The net log2 fold down-regulation of PLD in *M. phaseolina* inoculated Tx7000 was −3.4 while that of SC599 was −2.4. Therefore, the PLD pathway may not contribute to enhanced PA synthesis in both genotypes after *M. phaseolina* infection. The significantly lower PI(34:2) and DAG (34:2) content of *M. phaseolina-*inoculated SC599 indicated their contribution to enhanced PA content under pathogen infection through the PLC/DAGK pathway. The significantly up-regulated PLC gene (*Sb09g002320*) and non-significantly differentially expressed DAGK genes in SC599 bolsters this observation. The contribution of PLC/DAGK pathway to increased PA biosynthesis under *M. phaseolina* inoculation agree with previous reports that, pathogenic elicitors, in general, activate the PLC/DAGK pathway (de Jong *et al*., 2004; Laxalt & Munnik, 2002; Van der Luit *et al*., 2000; Den Hartog *et al*., 2003; Yamaguchi *et al*., 2003). On the other hand, the significantly higher PI(34:2), PI(36:4), and PI(36:2) content in inoculated Tx7000 is in agreement with the observed down-regulation of PLC genes. Therefore, although many DAGK genes are significantly up-regulated in Tx7000, the amount of PA generated through the PLC/DAGK pathway is comparatively lower compared to that of SC599 after pathogen infection.

Although PA plays an important role in plant defense against pathogens (Laxalt & Munnik, 2002; Munnik, 2001; Wang, 2004), the increased PA level could contribute to membrane bilayer destabilization, that results in membrane fusion and cell death (Welti *et al*., 2002). Moreover, a PA hike is likely to be upstream of an oxidative burst while exogenously applied PA can induce a partial oxidative burst in plants (de Jong *et al*., 2004). These studies suggest the cons of excessive PA to normal cellular function. The higher total PA content in Tx7000 compared to SC599 (across 7 and 10 DPI and treatments) suggested that Tx7000 may be more vulnerable to PA-hike-associated cell function retardation.

### Implications of galactolipids to phospholipid ratio and charcoal rot disease reaction

Plants synthesize additional galactolipids to replace phospholipids under phosphate deprivation, which results in an increased galactolipid:phospholipid ratio (Andersson *et al*., 2003; Hartel *et al*., 2000). However, low temperature stress increases the proportion of phospholipids and results in a decreased galactolipid:phospholipid ratio (Li *et al*., 2008; Uemura *et al*., 1995). In the current study, we observed a significantly greater galactolipid:phospholipid ratio in Tx7000 after *M. phaseolina* inoculation at 4 DPI. It may be possible that Tx7000 undergoes phosphate deprivation at the initial stages of pathogen infection. Interestingly, the exact opposite phenomenon was observed across 7 and 10 DPI. This indicates that the potential phosphate deprivation experienced by Tx7000 at the initial stages of infection would only be transient. The potential relationship between the galactolipid:phospholipid ratio and charcoal rot disease reaction deserves further investigation.

## ACKNOWLEDGEMENTS

The Kansas Grain Sorghum Commission is gratefully acknowledged for financial support of this research. The RNA sequencing project described in this work was carried out at the University of Kansas Medical Center while the lipidome analysis was performed at the Kansas Lipidomics Research Center Analytical Laboratory; instrument acquisition and lipidomics method development was supported by the National Science Foundation (EPS 0236913, MCB 0920663, MCB 1413036, DBI 0521587, DBI 1228622), Kansas Technology Enterprise Corporation, K-IDeA Networks of Biomedical Research Excellence (INBRE) of the National Institute of Health (P20GM103418) and Kansas State University. This paper is Contribution No. 20-###-J from the Kansas Agricultural Experiment Station, Manhattan.

## Supporting Information

**Table S1.** The lipid species (132) that were qualified for the analysis of variance (ANOVA), based on the limit of detection (>0.002 nmol) and coefficient of variation (< 0.3) and the mean concentrations (normalized intensity (%) per mg stalk dry weight) of 132 lipid species for SC599 and Tx7000 sorghum genotypes after receiving *M. phaseolina* (MP) and mock-inoculated control (CON) treatments at 4, 7, and 10 days post-inoculation (DPI).

**Table S2.** Significantly (q < 0.05) differentially expressed genes (related to lipid associated metabolic pathways) between SC599 (charcoal-rot-resistant) and Tx7000 (charcoal-rot-susceptible) sorghum genotypes in response to *Macrophomina phaseolina* inoculation at 7 days post-inoculation.

